# JAK Inhibitor Withdrawal Causes a Transient Proinflammatory Cascade: A Potential Mechanism for Major Adverse Cardiac Events

**DOI:** 10.1101/2024.09.25.615051

**Authors:** Ilya Gurevic, Loic Meudec, Xavier Mariette, Gaetane Nocturne, Sara S. McCoy

## Abstract

**Objective:** Our objective was to define the effect of JAK1/2 inhibitor (JAKinib) withdrawal on JAK/STAT biochemical response in the context of systemic rheumatic diseases.

**Methods:** We tested Type I (bind kinase active conformation) and Type II (bind kinase inactive conformation) JAKinibs in vitro using mesenchymal stromal cells and endothelial cells. We translated our findings in vivo studying NK cells from rheumatoid arthritis (RA) patients treated with Type I JAKinibs or methotrexate.

**Results:** Type I JAKinibs (ruxolitinib and baricitinib) increased phosphoJAK1 (pJAK1) and pJAK2 of IFNγ-stimulated MSCs and HUVECs in a time- and dose-dependent manner, with effect peaking after 24 hours. As expected, pSTAT1 was completely suppressed by JAKinibs. We found a marked and rapid *increase* of pSTATs upon discontinuation of Type I JAKinibs, that occurred to a lesser extent after Type II JAKinib withdrawal. Type I JAKinib withdrawal increased interferon and urokinase expression when compared to Type II JAKinib withdrawal. We found NK cells from RA patients taking Type I JAKinibs had a pro-inflammatory profile after JAKinib withdrawal compared to patients on methotrexate.

**Conclusions:** Type I JAKinibs paradoxically accumulate functionally defective pJAK. Upon withdrawal, the primed pJAKs are de-repressed and initiate a pSTAT signaling cascade, resulting in high interferon and urokinase. Type II JAKinibs do not cause pJAK accumulation, pSTAT cascade, and subsequent pro-inflammatory transcripts. The resultant cytokines and proteins produced from this cascade might be associated with adverse cardiac outcomes. Thus, JAKinib withdrawal is a possible mechanism contributing to the major adverse cardiac events described with JAKinib therapy.

## INTRODUCTION

JAK inhibitors (JAKinibs) are increasingly used to treat autoimmune diseases such as rheumatoid arthritis (RA) and psoriatic arthritis, among others. JAKinibs work by binding the adenosine triphosphate-binding site of JAK enzymes, which prevents them from carrying out their enzymatic activity. JAKinibs are generally grouped into two main classifications: Type I or Type II inhibitors. The licensed JAKinibs for RA treatment (e.g. baricitinib, filgotinib, tofacitinib, and upadacitinib) are all Type I inhibitors. Type I inhibitors interact with and stabilize the active conformation of JAK, leading to the abrogation of signaling through the JAK-STAT pathway. However, ubiquitination and degradation of JAK are also blocked leading to paradoxical accumulation of phosphorylation of the JAK activation loop (1). In contrast, Type II JAKinibs stabilize the inactive kinase conformation of JAK and are not associated with activation-loop phosphorylation when bound (2). No Type II JAkinibs are commercially available. Deucravacitinib, a TYK2 inhibitor, has a slightly different mechanism of action and uses allosteric inhibition of the enzymatic site of TYK2. Furthermore, it contains deuterium which increases the drug half-life and reduces unwanted metabolites (3).

JAKinibs, such as tofacitinib seem to be associated with a 33% increased risk of major adverse cardiac events (MACE) (4). MACE includes death from cardiovascular causes, nonfatal myocardial infarction, or nonfatal stroke. The risk of MACE is greatest among individuals ≥ 65 years old and with a past medical history of atherosclerosis (5). The European Medicines Agency safety committee has recommended restricted JAKinib use, though clear mechanistic explanation does not exist for why JAKinibs might increase cardiovascular risks (6).

Previous publications link Type I JAKinib withdrawal with life-threatening cytokine-rebound syndrome using cell lines to model myelofibrosis (another condition treated with JAKinibs) (1). The Type I JAKinib, ruxolitinib, prevents dephosphorylation of JAK2. In contrast, the Type II JAKinib CHZ868, did not cause JAK2 hyperphosphorylation (1). Type I JAKinibs increase loop phosphorylation that caused pathogenic signaling upon drug withdrawal, particularly in JAK2 V617F myelofibrosis patients (1).

Although this hypothesis was tested in myelofibrosis models using hematopoietic neoplasm and leukemia megakarytoblastic cells, the relevance of these findings to JAKinibs in the context of rheumatic disease is unknown. Our objective was to understand the downstream consequences of JAKinib withdrawal and how these might contribute to the possible MACE risk reported in rheumatology patients.

## METHODS

### Patients

All the work described was performed under the University of Wisconsin Health Sciences IRB 2018-0815 and Comité d’éthique de la recherche, CER Paris-Saclay 2020-A00509-30. Samples were collected from 01/2021 to 08/2021. Mesenchymal stromal cells (MSCs, fibroblasts) were generated from salivary gland tissue and express CD90 with a transcriptional fingerprint comprising *Pdgfra* and *Col15a1* (7). These cells are adherent to plastic and display the expected morphology in vitro (8, 9). Glands were derived from Sjögren’s disease (met 2016 ACR/EULAR criteria (10)) or control subjects (dry symptoms but no diagnosed autoimmune disease). Blood samples were collected from RA patients treated with methotrexate (MTX) or JAKinibs, referred to the Department of Rheumatology of Bicêtre Hospital (Université Paris-Saclay) during their standard of care. All patients fulfilled the ACR/EULAR 2010 criteria. Demographics are shown in Supplemental Tables S1 and S2.

### In Vitro Studies

We isolated, expanded, and froze MSCs from whole labial salivary glands as described previously (8, 9). For the described experiments, MSCs were expanded in α-minimal essential media (αMEM) (Corning, Tewksbury, MA) and 10% fetal bovine serum (FBS) (Sigma-Aldrich, St. Louis, MO) supplemented with 1% penicillin/streptomycin (Lonza, Walkersville, MD) and L-glutamine (Corning). Primary pooled human umbilical vein endothelial cells (HUVECs) (ATCC, Manassas, VA) were cultured at starting seeding density of 3,000-5,000 cells/cm^2^ in accordance with the manufacturer’s recommendations using vascular cell basal medium supplemented with bovine brain extract (ATCC). Optimal concentrations of IFNγ and drugs were determined in dose-finding experiments. MSCs and HUVECs were treated with the following reagents in the described experiments: IFNγ (10 ng/mL), ruxolitinib (1 µM unless otherwise specified), baricitinib (1 µM unless otherwise specified), CHZ868 (1 µM unless otherwise specified). Next, they were washed with PBS and treated with regular growth media with 0.01% DMSO, 10 ng/mL final concentration IFNγ+0.01% DMSO, or 10 ng/mL IFNγ + JAKinib for 48 hours. In the event of withdrawal, the cells were washed twice and then treated with our usual growth media with either 0.01% DMSO or with 10 ng/mL IFNγ+0.01% DMSO for a variety of time periods before harvest. In all cases the harvest of the cells was performed by collecting the conditioned media, washing once with PBS on ice, removing PBS and conducting on-plate lysis by scraping the cells from each 10 cm plate into 600 μL of 1x lysis buffer (cat. #9803, Cell Signaling Technology, Danvers, MA) supplemented with 1 mM EDTA, 1 mM PMSF, 1 mM NaF and 1 mM Na_3_VO_4_.

### Western Blot

After running the gel and transferring to a membrane, the membranes were blocked with 5% non-fat dry milk (cat. #sc-2325, Santa Cruz Biotechnology, Dallas, TX) in TBST (for phospho and total TYK2, 3% BSA in TBST was used per manufacturer instructions) at room temperature for ∼ 1 hr. Primary antibodies were diluted 1:1,000 in either 5% milk in TBST+0.02% sodium azide or 5% BSA in TBST+0.02% sodium azide, and their incubation with membranes was performed overnight at 4° C. After washing and incubation with HRP-conjugated secondary antibodies raised in goat against primary antibodies from rabbit or mouse (cat. #A120-101P, and #A90-116P, respectively, Bethyl Laboratories, Montgomery, TX) that were diluted 1:5,000 in 5% milk in TBST, membranes were imaged with Clarity™ Western ECL Substrate (cat.# 1705060, BioRad, Hercules, CA). Images were taken on an Amersham ImageQuant 800 (Cytiva, Marlborough, MA). Densitometric analysis was performed using ImageStudio Lite software (Li-Cor, Lincoln, NE).

Primary antibodies used were principally from Cell Signaling Technology: phospho-Akt (#4060), total Akt (#9272), phospho-Erk (#4370), total Erk (#9102), phospho-JAK1 (#3331), total JAK1 (#3332), phospho-JAK2 (#3771), total JAK2 (#3230), phospho-STAT1 (cat. # 9167), total STAT1 (cat. #9172), phospho-STAT2 (#88410), phospho-STAT3 (#9145), total STAT3 (#9139), total STAT4 (#2653), phospho-STAT5 (#9359), total STAT5 (#95205); others were from Abcam: total STAT2 (#32367), phospho-STAT4 (#ab313630); Bioss: phospho-TYK2 (#bs3437R), total TYK2 (#bs6662R); and Santa Cruz Biotechnology: GAPDH (#sc-25778).

### RNA Seq

We grew cells as described above and treated them as indicated with IFNγ/JAKinibs. After trypsinization, the cells were frozen and shipped to MedGenome, Inc. (Foster City, CA) for RNA isolation, reverse transcription, library construction and sequencing (Supplemental Methods).

### Real time quantitative PCR

After harvesting the MSCs in each treatment condition in TRIzol^®^, we used the Direct-zol RNA Miniprep columns for RNA isolation (cat. # Zymo Research, Irvine, CA). We generated cDNA using SuperScript IV reverse transcriptase (cat. #18090050, Invitrogen, Waltham, MA) per the manufacturer’s recommendations. qPCR was performed using the QuantiNova SYBR Green kit (*PLAU, PLAT*) (cat. # 208052, Qiagen, Germantown, MD) or the qPCR 2x Green Master Mix (*IFIT1*, *MX1*, *MX2* and *TNFSF15*) (cat. # 42-116PG, Apex Reagents and Chemicals). Primers sequences are shown in Supplemental Table S3. ΔΔC_t_ values (ΔΔC_t_=ΔC_t,IFNγ all the way_-ΔC_t,__) are plotted in the figure, and ΔC_t_ values were used for determining statistical significance.

### ELISA

Enzyme-linked immunosorbent assay (ELISA) was performed on conditioned media (CM) collected above the adherent cells. The uPA ELISA kits were performed per the manufacturer’s instructions (for the MSC experiment, cat. #IHUUPAKTT, Innovative Research, Novi, MI; for the HUVEC experiment, cat. #CSB-E04751h, CUSABIO, Houston, TX).

### Crosslinking assays

PBMCs from RA patients were collected the day of sampling and cultured overnight in RPMI with the addition of 10% FBS, 1% sodium pyruvate, 1% Hepes, 1% MEM, 1% penicillin/streptomycin and 200 UI/mL of IL-2.The day after, PBMCs were stimulated by crosslinking as previously described (11). Briefly, PBMCs were seeded on 96-well plates (Maxisorb, Thermofisher Scientific) previously coated with anti-CD16 (cat# 555404, BD Bioscience) or isotype control and incubated at 37°C 5% CO2 for five hours with IL-2 (10 UI/mL). Golgistop was added after 1h. PBMCs were then washed and stained for FACS analysis. IFNγ and TNFα were assessed after permeabilization.

### FACS analysis

Flow cytometry was performed on FACS Canto II (Beckman Coulter) and analysis made using FlowJo software. NK cells were gated as CD56+CD3-among living cells. Results were expressed as a percentage of positive cells for each marker. The antibodies used are indicated in Supplemental Table S4.

### Statistical Analysis

ΔΔC_t_ statistical significance was computed by two-sample unpaired Student’s t-test on ΔC_t_ values. Elsewhere, multiple comparison statistical significance was determined via a mixed-effects parametric model with multiple-comparison testing using Bonferroni’s correction. Statistical analysis was carried out by GraphPad Prism software (Graphad, Software, La Jolla, CA).

## RESULTS

### The Type I JAKinib, ruxolitinib, promotes pJAK1 and pJAK2 phosphorylation

Ruxolitinib increased IFNγ-induced phosphoJAK1 (pJAK1) and pJAK2 relative to total JAK1 and JAK2, respectively. Peak phosphorylation of JAK1 and JAK2 occurred around 24 hours but persisted at least 72 hours (Figure 1A-B). As expected, ruxolitinib quickly suppressed downstream pSTAT1 levels (Figure 1C). There were no substantive increases in pSTAT2-5, or pTYK2 with the addition of ruxolitinib to IFNγ; however, there was a subtle increase in C-jun and cyclin D1 1-2 hours after the addition of ruxolitinib (Supplemental Figure S1).

**Figure 1.**
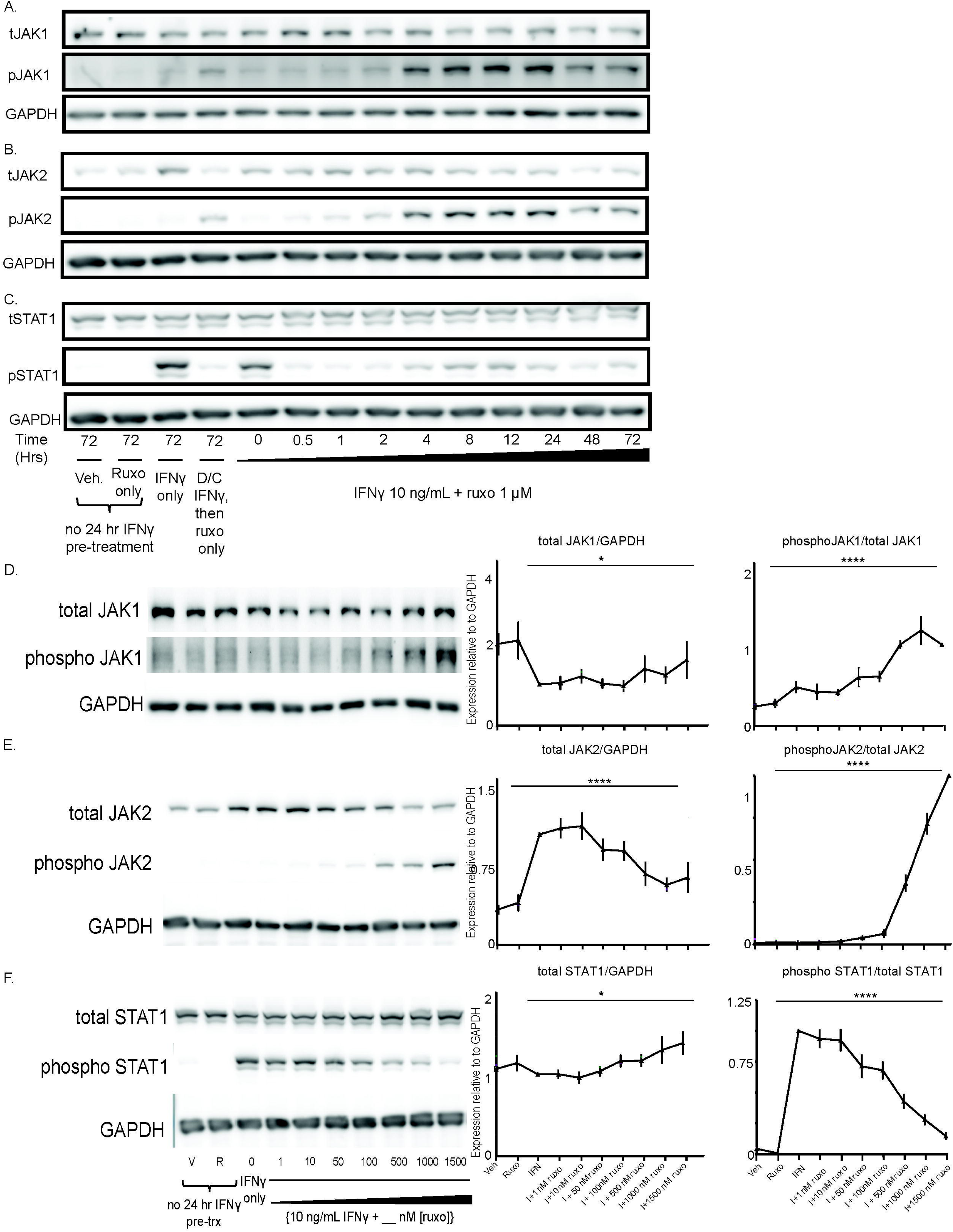
The Type I JAKinib, ruxolitinib, promotes pJAK1 and pJAK2 phosphorylation. We treated MSCs with vehicle, ruxolitinib (1 µM), IFNγ only (10 ng/mL), IFNγ then switching to ruxolitinib only, or IFNγ 10 ng/mL in combination with ruxolitnib 1 uM for varying periods of time or for 48 hours at varying doses of ruxolitinib (0-1500nM). At each time or dose of ruxolitinib, we harvested cells for protein and performed western blot for the indicated target. A) Total JAK1 is stable regardless of treatment time with ruxolitinib but the proportion of pJAK1 increases starting at four hours and peaking after 24 hours of treatment; B) the proportion of pJAK2 increases with ruxolitinib treatment, starting at four hours and peaking after 24 hours; C) pSTAT1 is suppressed almost immediately after introduction of ruxolitinib. Fig 1D-F were performed with replicates (n=6; 3 SjD and 3 control) of our treatment conditions on the right with a representative western on the left; D) pJAK1 proportion increases in a dose-dependent manner with increasing ruxolitnib; E) pJAK2 proportion increases as ruxolitinib dose increases; F) pSTAT1 decreases in a dose response relationship with increasing ruxolitinib. Error bars represent standard deviation; *p=0.05; p=****<0.0001 represent the ANOVA-derived significance for the difference between all values.

We found that increasing doses of ruxolitinib decreased total JAK1 and total JAK2 with maximal reduction around 500 nM ruxolitinib (Figure 1D and 1E). Phosphorylation of JAK1 and JAK2 relative to total JAK increased in a dose-dependent fashion starting at around 50nM of ruxolitinib (Figure 1D-E). pSTAT1 reached negligible levels at 1500 nM ruxolitinib (Figure 1F). There were no differences in JAK/STAT phosphorylation patterns between SjD and control MSCs (data not shown).

Ruxolitinib alone, without the presence of IFNγ, did not increase the expression of any of our tested pathways. These results show the addition of ruxolitinib to IFNγ increases pJAK1 and pJAK2 phosphorylation, confirming ruxolitinib is a Type I JAKinib capable of promoting phosphorylation of JAK1/2. We also found that in the IFNγ alone treatment did not substantially increase pJAK1 and pJAK2 compared to withdrawal conditions; however, pJAK1 and pJAK2 were greater in the IFNγ alone treatment condition than vehicle or ruxolitinib alone (Supplemental Figure S2).

### While Type I JAKinibs promote JAK phosphorylation, Type II JAKinibs do not

We replicated our initial results using increasing concentrations of bariticinib, another Type I JAKinib. We found that baricitinib increased pJAK2 relative to total JAK2 in a dose-dependent manner (Figure 2A), similar to ruxolitinib. CHZ868 is a Type II JAK inhibitor that interacts with JAK2 in the inactive conformation. We showed that CHZ868 did not increase pJAK2 (Figure 2C). As expected, both baricitinib and CHZ868 decreased pSTAT1 in a dose dependent manner (Figure 2C-D). Viability was ascertained by cell morphology with no significant effects of JAKinib dose on cell viability. 1000 nM JAKinib was selected for the remainder of the experiments because it was lowest concentration associated with clear reduction of pSTAT1.

**Figure 2.**
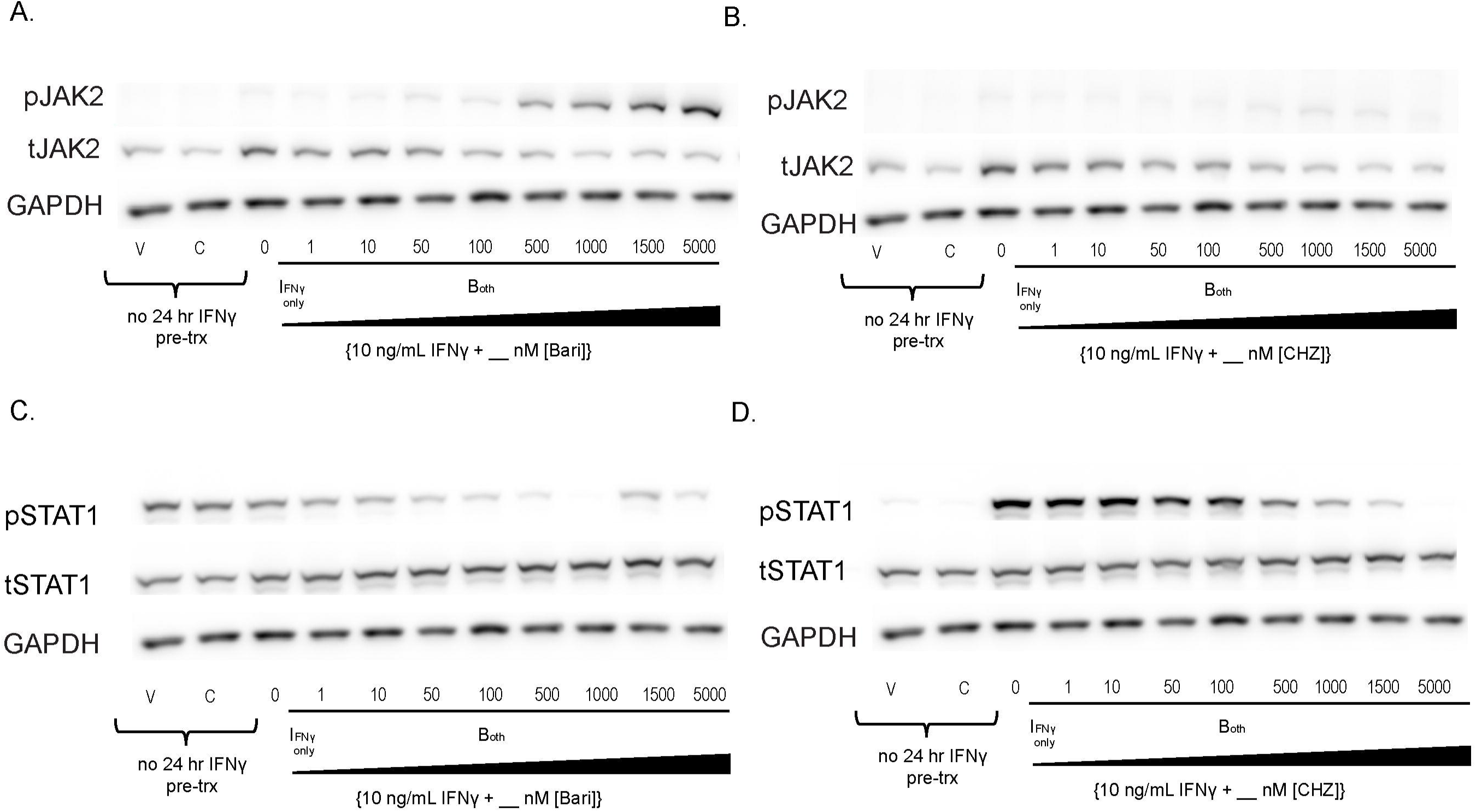
Type I JAKinibs promote JAK phosphorylation while Type II JAKinibs do not. We treated SG-MSCs with vehicle, JAKinib alone (C=CHZ868; B=baricitinib), or varying doses of baricitinib or CHZ868 with IFNγ (10 ng/mL). After 48 hours, the cells were harvested, and protein was isolated for western blot to the shown targets. A) There is a slight decrease in total JAK2 with increasing barictinib but a more pronounced increase in pJAK2 relative to total JAK2; B) total JAK2 decreases with increasing CHZ868 but there is no pJAK2 phosphorylation relative to total JAK; C) increasing concentrations of baricitinib does not affect total STAT1 but drives a dose-related reduction in pSTAT1; D) increasing concentrations of CHZ868 does not affect total STAT1 but drives a dose-related reduction in pSTAT1.

### Type I JAKinib withdrawal causes a transient surge in pSTAT, Type II JAKinib withdrawal does not

After serum starvation, we treated MSCs with IFNγ for 24 hours, washed the cells, and treated them with IFNγ plus a JAKinib (ruxolitinib, baricitinib, or CHZ868) for 48 hours, washed the cells, and withdrew the JAKinib from each condition while continuing IFNγ. We found that withdrawal of ruxolitinib and baricitinib led to decreased pJAK2 over three hours (Figure 3A). In contrast CHZ868, lacking increased pJAK2 at baseline, had near undetectable levels of pJAK2 (Figure 3A). Withdrawal of ruxolitinib and baricitinib led to marked increase in pSTAT1 (Figure 3B). In contrast, CHZ868 led to a slight increase in pSTAT1 (Figure 3B). Interestingly, the expression of pSTAT1 appeared greater with ruxolitinib and baricitinib withdrawal than de novo treatment of MSCs with 10 ng/mL IFNγ. We tested pSTAT2-5 and found that hyperphosphorylation of STAT on Type II JAK inhibitor withdrawal was most pronounced with STAT2 but also occurred in STAT3 and 5 (Figure 3C-F). Because IFN also can induce JAK to activate the MAPK and PI3K/AKT systems, we tested whether the withdrawal hyperphosphorylation phenomenon affects other related pathways (12). We found that although both phospho-ERK and phospo-AkT increased after withdrawal, there was no clear association with either a Type I or Type II JAKinib-specific response (Supplemental Figure S3).

**Figure 3.**
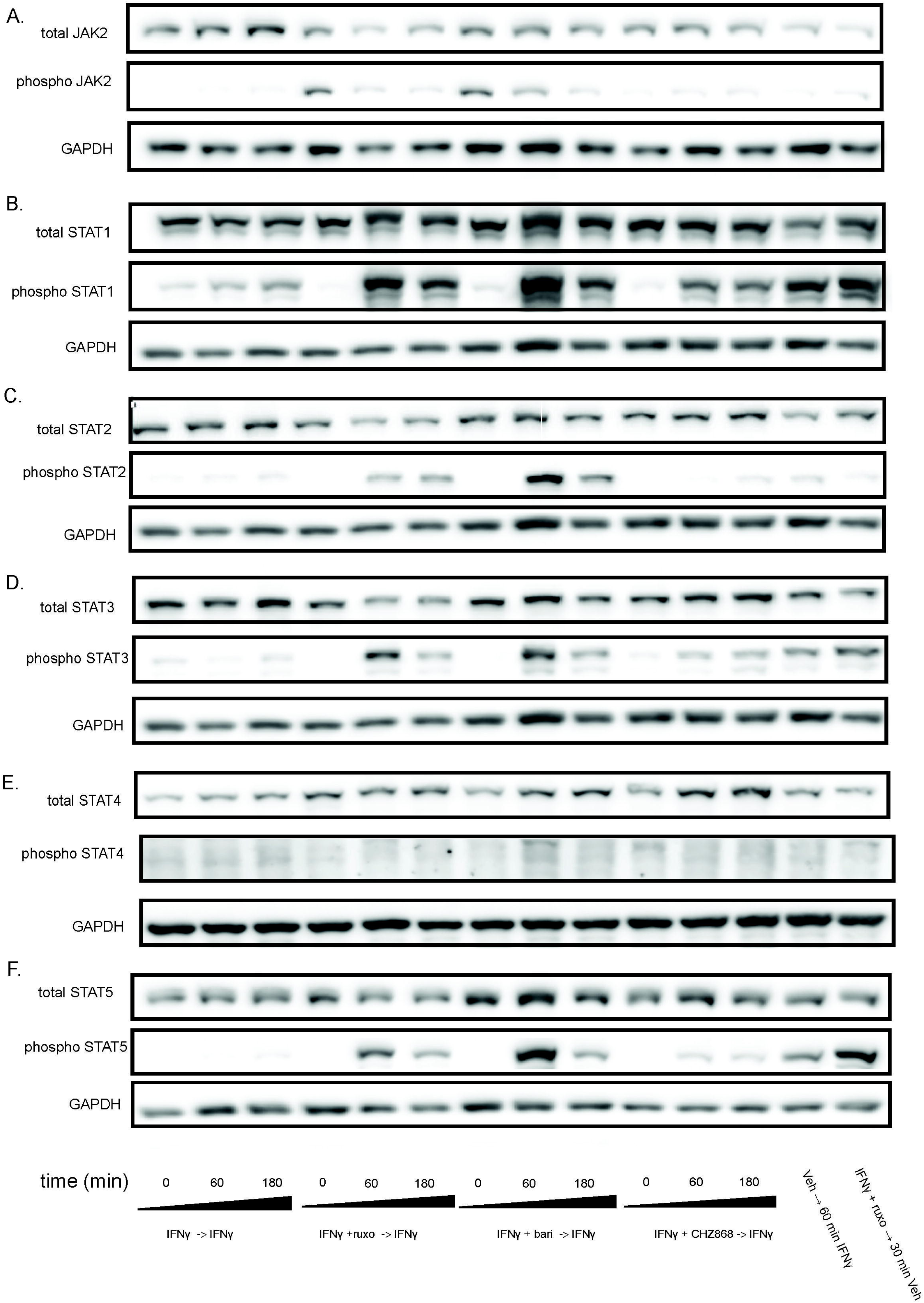
Withdrawal of Type I JAKinibs increased STAT phosphorylation, a phenomenon present but markedly lower in Type II JAKinibs. We treated MSCs with IFNγ 10ng/mL then added IFNγ alone, IFNγ and ruxolitinib (1 µM), IFNγ and barictinib (1 µM), or IFNγ and CHZ868 (1 µM). After 48 hours, we removed the medium, washed the cells, and added back media with only IFNγ. Cells were harvested at the indicated time points and western blot was performed on the resulting protein lysates. GAPDH was used as a control. A) pJAK2 peaks at time zero after exposure to baricitinib and ruxolitinib and then decreases. There is no increase in pJAK2 with CHZ868 treatment; B) pSTAT1 proportion peaks after 60 minutes and is greater after Type I JAKinib exposure than Type II JAKinib exposure; C) The pSTAT2 proportion peaks 60 minutes after type I JAKinib withdrawal but this phenomenon is not seen with Type II JAKi withdrawal; D) The pSTAT3 proportion peaks 60 minutes after type I JAKinib withdrawal but this phenomenon is seen to a much lesser extent with Type II JAKinib withdrawal; E) There is no clear effect on pSTAT4; F) The pSTAT5 proportion peaks 60 minutes after type I JAKinib withdrawal but this phenomenon is seen to a much lesser extent with Type II JAKinib withdrawal.

### JAK hyperphosphorylation and STAT phosphorylation after withdrawal of Type I JAKinibs occur in other cell types

We used MSCs as a rosetta stone for cellular response to JAKinib withdrawal, but needed to confirm whether this phenomenon was cell-specific or seen across various cell types. We found that, akin to MSCs, HUVECs show increased JAK2 phosphorylation with exposure to Type I JAK inhibitors but not Type II JAK inhibitors (Figure 4A). On JAKinib withdrawal, pSTAT1 increased more in Type I than Type II JAKinibs (Figure 4B). Finally, just as with the MSC response, HUVECs show marked reduction in phosphorylation of STAT2 and STAT5 on Type II JAKinib withdrawal compared to Type I JAKinib withdrawal (Figure 4C-D).

**Figure 4.**
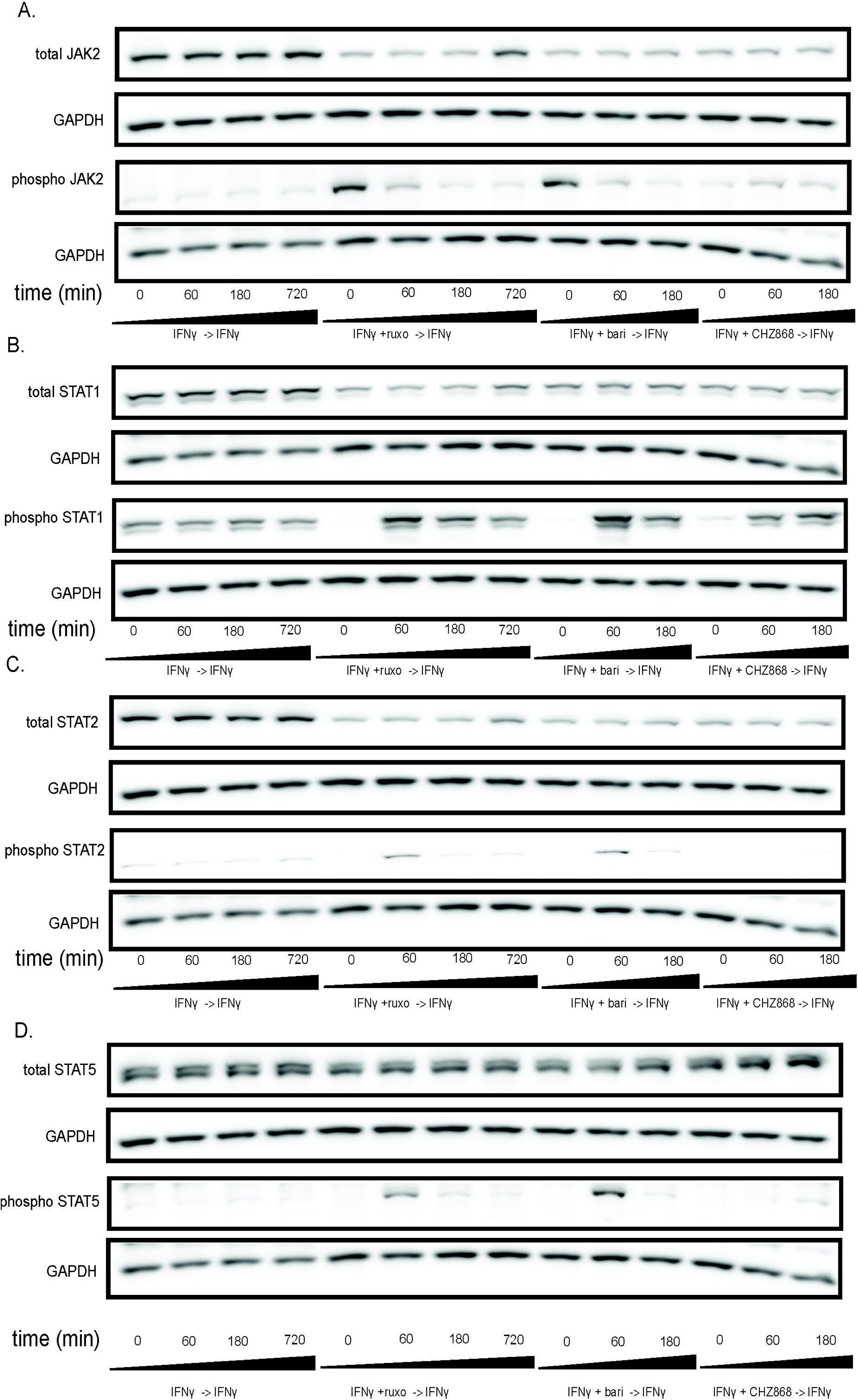
The JAKinib withdrawal phenomenon observed in MSCs can be extended to other cell types. We treated HUVECs with IFNγ for 24 hours then proceeded with the following conditions: IFNγ alone, IFNγ and ruxolitinib (1 µM), IFNγ and barictinib (1 µM) or IFNγ and CHZ868 (1 µM). After 48 hours, we removed the medium, washed the cells, and added back media with only IFNγ. Cells were harvested at the indicated time points and western blot was performed on the resulting protein lysates. GAPDH was used as a control. A) pJAK2 peaks at time zero after exposure to baricitinib and ruxolitinib and then decreases. There is no increase in pJAK2 with CHZ868 treatment; B) pSTAT1 proportion peaks after 60 minutes and is greater after Type I JAKinib exposure than Type II JAKinib exposure; C) The pSTAT2 proportion peaks 60 minutes after type I JAKinib withdrawal but this phenomenon is not seen with Type II JAKinib withdrawal; D) The pSTAT5 proportion peaks 60 minutes after type I JAKinib withdrawal but this phenomenon is seen to a much lesser extent with Type II JAKinib withdrawal.

### IFN and PLAU (urokinase) increases with Type I JAKinib withdrawal

To obtain a global view of the differential transcriptional pathways between Type I and II JAKinib withdrawal, we performed RNA sequencing comparing 3 hour withdrawal of ruxolitinib to 3 hour withdrawal of CHZ868. Our RNA sequencing results showed Type I interferon (IFN) signaling pathway transcripts were downregulated in CHZ868 withdrawal relative to ruxolitinib withdrawal conditions (Figure 5A). We tested IFN and related pathways with qPCR including IFIT1, MX1, MX2, and TNFSF15. MX2 transcripts were significantly higher after ruxolitinib withdrawal than CHZ868 withdrawal (Figure 5B). MX1, but not IFIT1 and TNFSF15 transcripts, are upregulated with ruxolitinib discontinuation compared to CHZ868 discontinuation but did not achieve statistical significance (Supplemental Figure S4 A-C)

**Figure 5.**
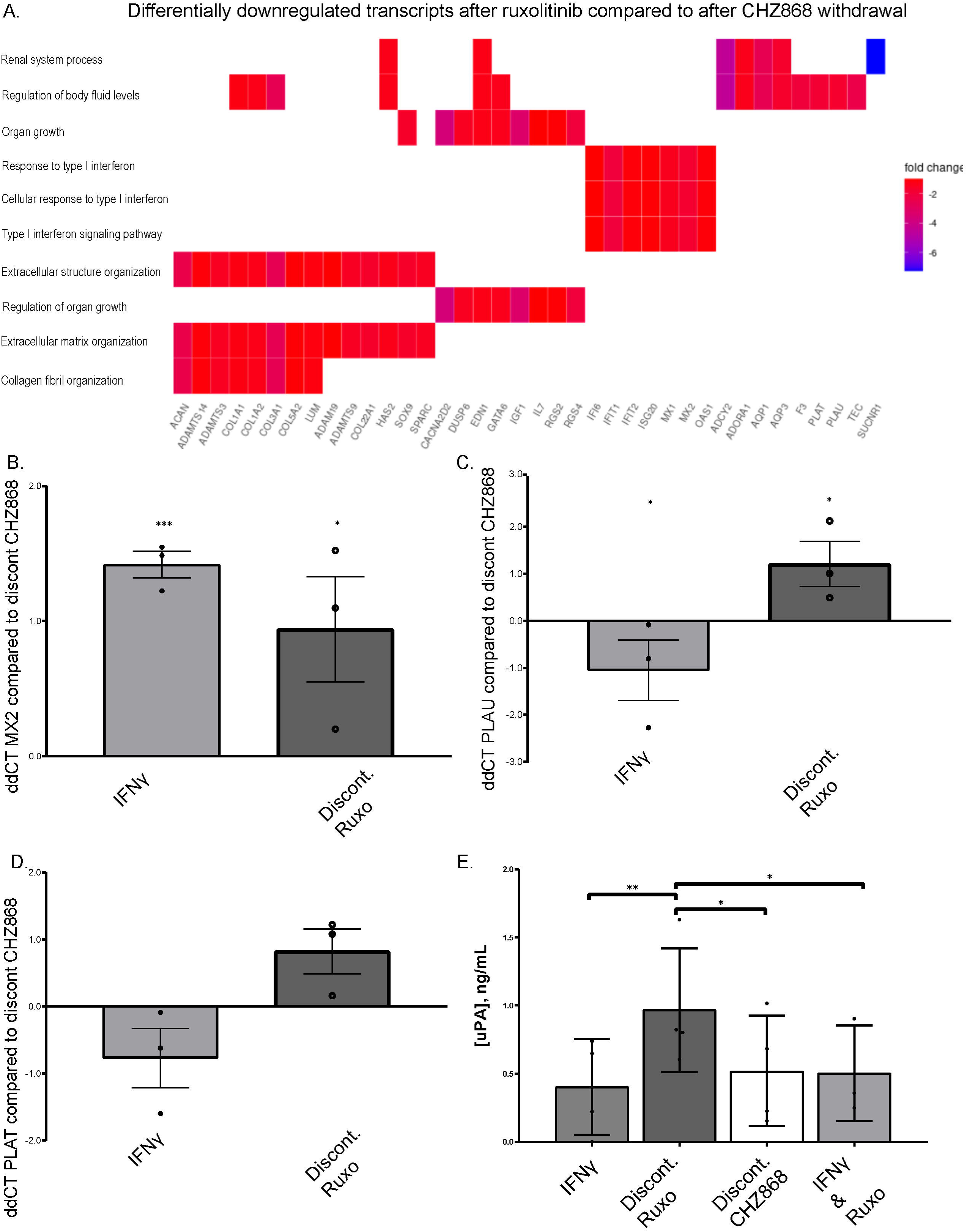
IFN and PLAU (urokinase) are increased after ruxolitinib compared to CHZ868 withdrawal. A) Bulk RNA seq was performed of MSCs that were exposed to the following treatments IFNγ 10 ng/mL and ruxolitinb (1000 nM), wash cells, then continue treatment only with IFNγ (ruxolitinib withdrawal) or IFNγ and CHZ868 (1000 nM) treatment, wash cells, then continue treatment with only IFNγ (CHZ868 withdrawal). The heatmap shows differentially downregulated transcripts in ruxolitinib compared to CHZ868 withdrawal; B) RT-qPCR of RNA isolated from MSCs after treatment in the above conditions and harvested 3 hours later show MX2 transcripts are significant higher after ruxolitinib discontinuation than after CHZ868 discontinuation. ΔΔC_t_ values (ΔΔC_t_=ΔC_t,DC CHZ_-ΔC_t,__) are plotted in the figure and statistical significance was computed by two-sample unpaired Student’s t-test on ΔC_t_ values compared to CHZ868 withdrawal conditions; C) qPCR of MSCs after treatment in the above conditions and harvested 3 hours later show PLAU transcripts are significantly higher after ruxolitinib discontinuation than after CHZ868 discontinuation. ΔΔC_t_ values (ΔΔC_t_=ΔC_t,DC CHZ_-ΔC_t,__) are plotted in the figure, and statistical significance was computed by two-sample unpaired Student’s t-test on ΔC_t_ values; D) qPCR of MSCs after treatment in the above conditions and harvested at 3 hours post-withdrawal show that PLAT transcripts show a trend toward being higher after discontinuation of ruxolitinib than after CHZ868 discontinuation; E) ELISA of conditioned medium aspirated from MSCs after 3 hours of above treatment conditions show that PLAU is significantly increased in the culture media of MSCs after ruxolitinib withdrawal compared to other conditions; statistical significance was determined via a mixed-effects parametric model with multiple-comparison testing using Bonferroni’s correction. *p<0.05; ** p<=0.01; ***p<0.001

We found that tissue plasminogen activator [PLAT; tPA] and urokinase-type plasminogen activator [PLAU; uPA] transcripts were downregulated in CHZ868 withdrawal relative to ruxolitinib withdrawal (Figure 5A; Supplemental Table S5) in our RNA sequencing results. Because PLAT and PLAU are also associated with atherosclerosis/thrombotic pathways and JAKinib therapy is associated with MACE, we focused on these PLAT and PLAU transcripts for further analyses. We confirmed PLAU transcripts increased more with ruxolitinib discontinuation than CHZ868 discontinuation by qPCR and saw a diminished but similar trend with PLAT (Figure 5C-D). We found that after ruxolitinib withdrawal, MSCs made significantly more uPA than after CHZ868 discontinuation (Figure 5E). Conditioned media assessment of uPA from HUVECs was below the level of detection (not shown), indicating that the STAT signaling cascade produced by JAKinib withdrawal might have cell-specific consequences.

### IFNγ and TNFα related inflammatory transcripts are increased after Type I JAKinib withdrawal

NK cells might contribute to autoimmunity through exposure of autoantigens, promoting maturation of dendritic cells and differentiation of monocytes (13, 14). Furthermore, NK cells are a major producer of IFNγ and are activated by Type I IFNs, linking their relevance in autoimmunity to our observed findings (15, 16). The JAK/STAT pathway is highly involved in these cells, making them a good model to study the effect of Type I JAKinib withdrawal in patients.

We analyzed the functionality of NK cells from RA patients treated with methotrexate, baricitinib, or tofacitinib after overnight drug withdrawal (Figure 6A). To do so, we stimulated PBMCs by crosslinking with anti-CD16 and assessed the degranulation marker CD107a and intracellular production of IFNγ and TNFα (Figure 6B).

**Figure 6.**
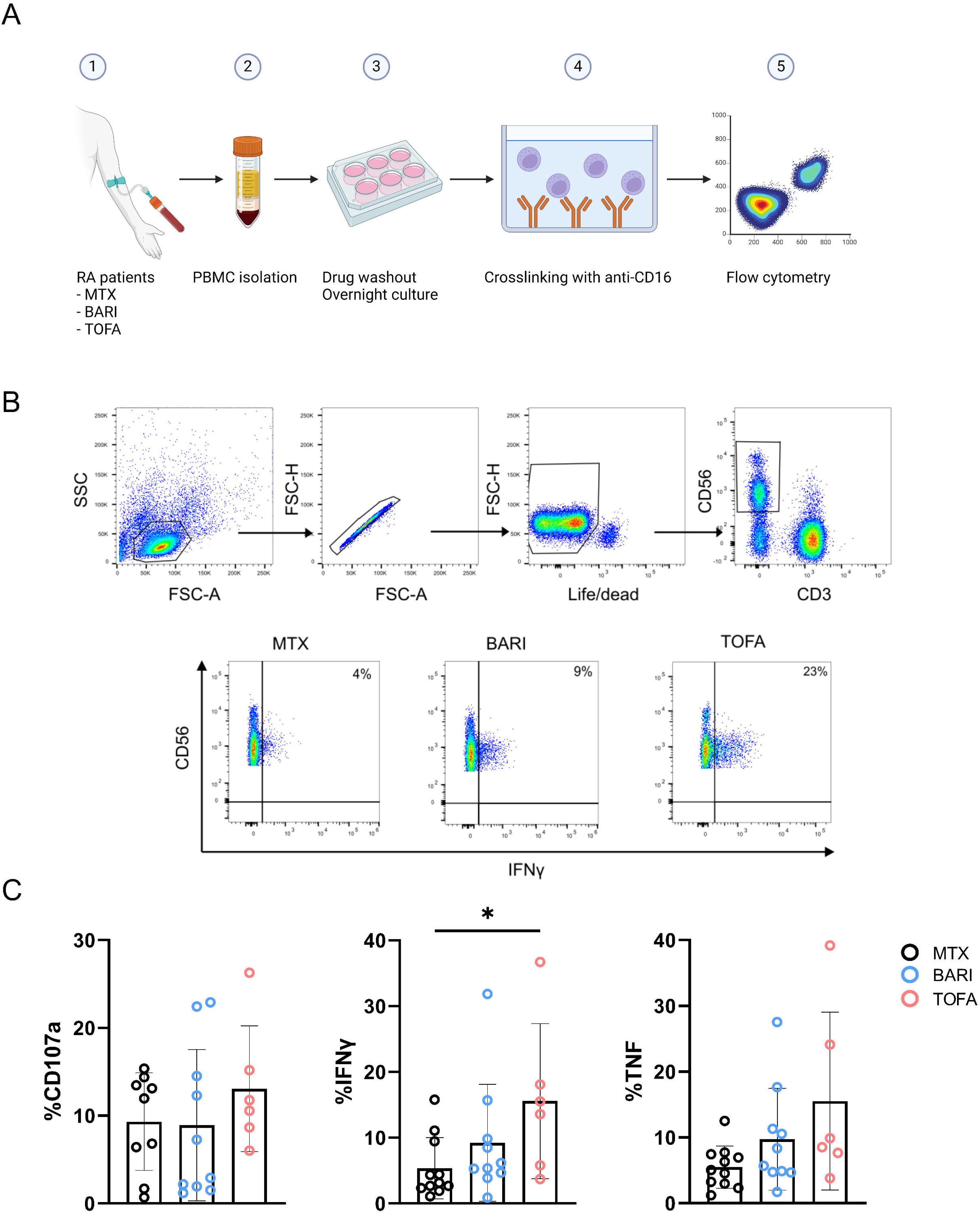
JAKinib withdrawal results in patient-derived NK cell activation. PBMCs from RA patients treated with JAKinibs (tofacitinib [TOFA] or baricitinib [BARI]) or methotrexate (MTX) were stimulated upon CD16 by crosslinking for 5 hours after overnight withdrawal of the drug. CD107a and intracellular IFNγ and TNFα were assessed by flow cytometry. A) Schematic for patient PBMC crosslinking; B) flow cytometry gating strategy. NK cells were defined as CD3-CD56+; C) CD107a, IFNγ and TNFα expression by flow cytometry. NK cells from patients treated with JAKinibs expressed more IFNγ and those treated with JAKinibs trended to greater NK cell activation overall. Kruskal-Wallis test was used. p<0.05 was considered significant.

We found that NK cells from patients taking the Type I JAKinib tofacitinib expressed significantly more IFNγ than those taking methotrexate. There was a trend toward higher CD107a and TNFα expression in NK cells from patients taking Type I JAKinibs compared to methotrexate. These findings support that upon withdrawal from JAKinibs, NK cells demonstrate a pro-inflammatory profile.

## DISCUSSION

We describe a mechanism linking Type I JAKinib exposure to a possible adverse outcome associated with their use, MACE (Figure 7). Upon in vitro IFNγ exposure, Type I JAKinibs increase phosphorylation of JAKs compared to Type II JAKinibs; however, both similarly suppress downstream signaling pathways. We revealed a divergent response to JAKinib withdrawal: Type I JAKinib withdrawal results in a downstream STAT phosphorylation surge relative to Type II JAKinib withdrawal. We showed differential Type I and Type II JAKinib withdrawal signaling cascades results in greater IFN- and coagulation cascade-related transcription and protein production. More specifically, we found that in a cell-specific manner, Type I JAKinib withdrawal promotes pro-inflammatory NK cells. Type I JAKinib withdrawal also stimulates uPA production from stromal cells.

**Figure 7.**
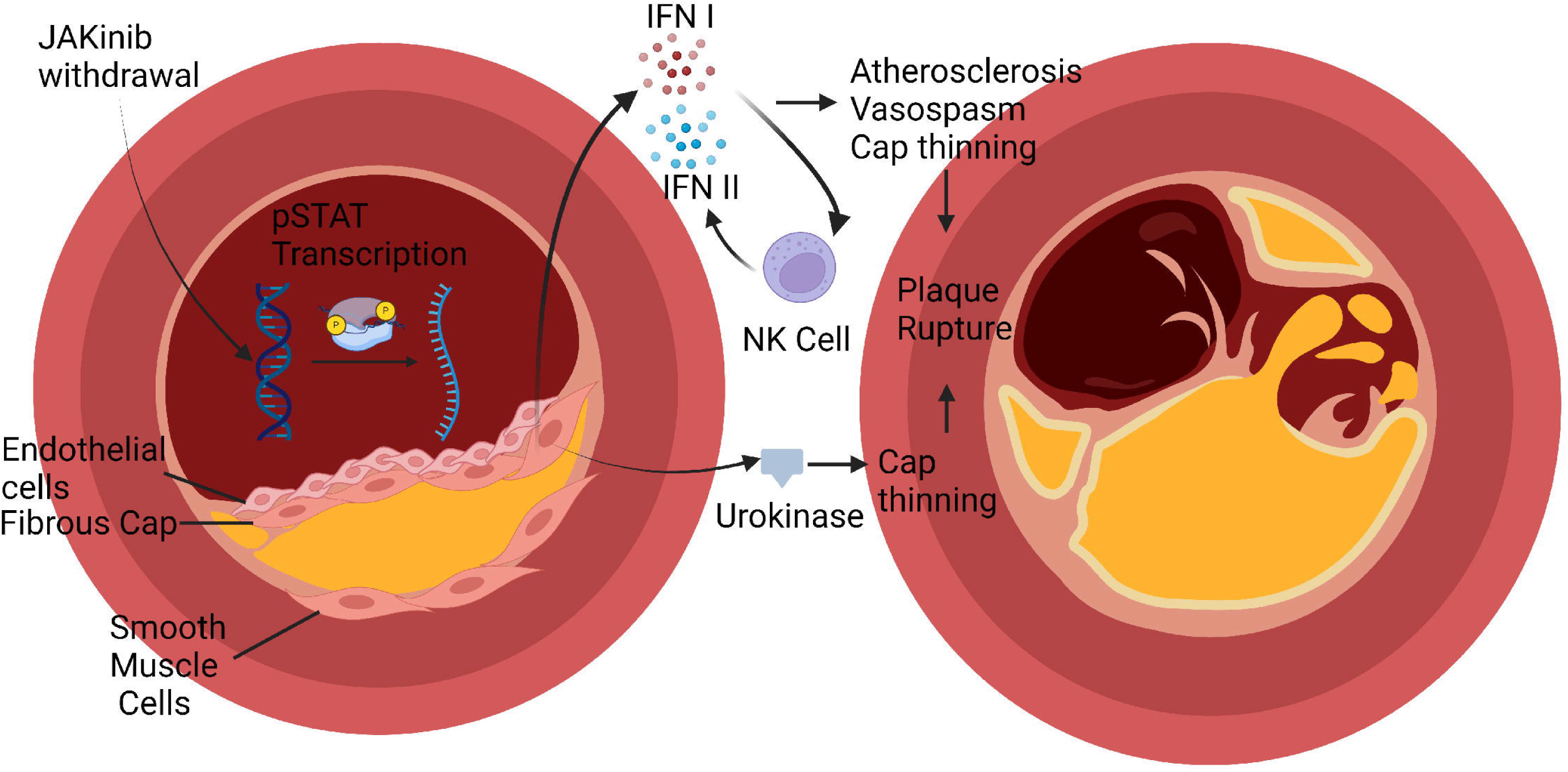
Graphical Abstract. In rheumatic disease patients, JAKinib withdrawal could drive a pro-inflammatory cascade that might contribute to major cardiac adverse events. Type I JAKinib exposure drives JAK hyperphosphorylation that is de-repressed upon JAKinib withdrawal. The subsequent spike in pSTAT signaling results in Type I IFN and urokinase production. Type I IFN activates NK cells toward a pro-inflammatory phenotype. Ultimately, JAKinib withdrawal might contribute to atherosclerosis and plaque rupture through IFN-mediated atherosclerosis, vasospasm, and fibrous cap thinning.

JAKinibs, such as tofacitinib, are associated with a 33% increased risk of MACE among patients more than 50 years old with at least one cardiovascular risk factor (4). Cardiovascular risk factors include a history of coronary artery disease, cerebrovascular disease or peripheral artery disease (5). Mechanisms driving this finding remain unclear. We made two relevant findings to possibly explain increased MACE with JAKinib use. The first finding was that Type I JAKinib withdrawal resulted in NK cells with a proinflammatory phenotype compared to patients on methotrexate. Interestingly, this result is the opposite of what is observed when exposing *in vitro* NK cells to JAKinib without withdrawal, leading to reduced functionality and activation profile (11). It is possible that Type I IFN generated from stromal cells upon JAKinib withdrawal potentiates other cells, such as NK cells, to produce Type II IFN. This might ultimately lead to IFN-driven effects on atherosclerosis and cardiovascular disease. Multiple studies have linked IFN treatment to adverse cardiac events (17–20). These studies also indicate that the most likely risk factor for cardiotoxicity with IFN exposure was previous cardiac disease (17). This effect is theorized to be caused by increased oxygen demand from the flu like symptoms generated by IFN exposure leading to infarction or arrhythmia. IFN might also potentiate atherosclerosis through several mechanisms including supporting foam cells, endothelial dysfunction, local immune cell activation, and NET formation, among other pathways (21). In addition to IFN treatment as a risk factor for cardiac events, high IFN in diseases like systemic lupus erythematosus, might be associated with MACE (22). Thus, it is possible that increased IFN produced upon JAKinib withdrawal might contribute to the increased MACE seen with JAKinib use among patients with atherosclerotic disease. In patients using immediate release tofacitinib, whose half-life is three hours, it is possible that the serum level of the drug sometimes decreases below the efficient doses for inhibiting JAK inhibition. This could stimulate JAKinib withdrawal and transiently stimulate IFN secretion. This risk might be especially pronounced if patients miss or delay their dosing. If this is the case, the use of deuterated JAK inhibitors, which prolong JAKinib half-life, might reduce this potential adverse effect.

The second effect of Type I JAKinhib withdrawal that we appreciated was the increased production of uPA in stromal cells, important players in initiation and advancement of atherosclerotic disease. uPA converts plasminogen into plasmin. Plasmin is a serine protease with broad effects on extracellular matrix (ECM) including fibronectin, laminin, and vitronectin (23). At first glance, elevated uPA might seem protective against MACE, but research suggests uPA might promote plaque rupture. uPA is associated with atherosclerosis in human coronary artery plaque (24). Its receptor, urokinase-type plasminogen activator receptor, is highly expressed in macrophages in ruptured plaques (25). uPA overexpression in mice causes intraplaque hemorrhage and fibrous cap disruption, potentially through overexpression of MMPs (26). A recent elegant study sought to define the proteomics of mouse and human plaque rupture in parallel, hypothesizing that plaque rupture is started by artery wall extracellular matrix proteolysis (27). Transgenic mice that overexpressed macrophage urokinase had aortic tissue with inflammatory signaling molecules and protein involved in cell adhesion, cytoskeleton, and apoptosis. These mice had findings in their vessels that paralleled human plaque including decreased basement-membrane proteins. They theorized high local uPA drives proteases (e.g. plasmin, MMP9, MMP12) that cleave laminin, nidogen HSPG2, and Collagen XVIII. This results in a loss of basement membrane proteins and plaque rupture (27). More recently, PBMCs from patients with plaque rupture were analyzed by high throughput sequencing and revealed five major genes increased in PBMCs from patients with plaque rupture, two of these five genes were PLAU and PLAUR (28). The authors of this study posit that hub genes such as PLAU and PLAUR serve as biomarkers for prospective prediction of atherosclerotic plaque rupture (28). Finally, PLAU and PLAT are increased in carotid plaques and associated with unstable plaque (29). Thus, uPA elevation is relevant to plaque rupture and thrombosis and might explain a component of MACE seen with Type I JAKinib use.

If our hypothesis of a paradoxically pro-inflammatory effect of Type I JAKinib withdrawal is correct, it would be interesting to assess whether a lack of compliance, which would favor this rebound effect in clinical practice, could be associated with a higher cardiovascular risk in RA patients.

In conclusion, we identify novel and plausible mechanisms related to Type I JAKinib withdrawal that might explain the MACE seen with JAKinib therapy in certain populations relevant to rheumatology. These results might affect clinical practice; for example, some experts might reconsider recommendations to hold JAKinibs before prolonged travel or favor Type II or modified Type I JAKinibs. Future studies are required to further test the sequelae of the identified withdrawal phenomenon in vivo using animal models of disease.

## Data availability statement

All data relevant to this study are included in the article.

## Ethics approval

This study was approved by the University of Wisconsin Health Sciences IRB (IRB# 2018-1815) and Comité d’éthique de la recherche, CER Paris-Saclay 2020-A00509-30 and complies with the Declaration of Helsinki.

Presented at: Preliminary results from this study have been presented by McCoy S, *et al* at the 2022 and 2023 American College of Rheumatology meeting.

## Data availability statement

The data used to generate the findings presented herein are available upon request to the corresponding author.

## Acknowledgements

We thank Dr. Jacques Galipeau for his mentorship and editorial input. The authors would like to thank the University of Wisconsin Carbone Cancer Center for the use of its Translation Science shared services (supported in part by NIH/NCI P30 CA014520 to the UW Comprehensive Cancer Center Support) to complete this research.

