## Supplemental files for "JAK Inhibitor Withdrawal Causes a Transient Proinflammatory Cascade: A Potential Mechanism for Major Adverse Cardiac Events"

**Supplemental Table S1: Characteristics of the patients from which MSCs were derived**

| Age | Sex | SjD | Figure(s) |
| --- | --- | --- | --- |
| 39 | **F** | **control** | **5** |
| 60 | **F** | **SjD** | **1** |
| 52 | **F** | **control** | **1** |
| 52 | **F** | **control** | **5** |
| 51 | **F** | **control** | **1,5** |
| 30 | **F** | **SjD** | **1** |
| 31 | **F** | **SjD** | **1,2,3** |
| 31 | **F** | **control** | **1** |
| 49 | **F** | **control** | **1** |

SjD patients met 2016 ACR/EULAR criteria. Control subjects

were referred for labial salivary gland biopsy due to dryness

symptoms but did not have a defined autoimmune disease.

**Supplemental Table S2: Characteristics of the patients from which NK cells were derived**

|  | **MTX, n=9 (%)** | **JAKi, n=16 (%)** |
| --- | --- | --- |
| Age, mean ± SD | 65.2 ± 14.5 | 57.8 ± 16.6 |
| Women | 8 (88.9) | 14 (87.5) |
| **Rheumatoid arthritis** |  |  |
| Disease duration (years), mean ± SD | 4.2 ± 4.2 | 14.9 ± 8.4 |
| Erosive | 3/8 (37.5) | 11 (68.8) |
| Rheumatoid Factor positivity | 6/8 (75) | 9/15 (60) |
| ACPA positivity | 6/8 (75) | 11/15 (73.3) |
| DAS28, mean ± SD | 2.1 ± 1.6 | 3 ± 1.1 |
| **Treatments** |  |  |
| Corticosteroids | 4 (44.4) | 4 (25) |
| Average corticosteroids dose, mean ± SD | 7 ± 3.6 | 6.2 ± 2.5 |
| MTX | 9 (100) | 9 (56.2) |
| Baricitinib | NA | 10 (62.5) |
| Tofacitinib | NA | 6 (37.5) |
| Prior bDMARD | 0 | 16 (100) |

Data are n (%) unless otherwise indicated

| Supplemental Table S3. Primer sequences and targets | | |
| --- | --- | --- |
| Target | Forward | Reverse |
| β-actin | 5’-CATCACGATGCCAGTGGTACG-3’ | 5’-AACCGCGAGAAGATGACCCAG-3’ |
| IFIT1 | 5'-GCCCTGGAGTACTATGAGCGG-3' | 5'-GCTGATATCTGGGTGCCTAAGGAC-3' |
| MX1 | 5′-GGCTGTTTACCAGACTCCGACA-3′ | 5′-CACAAAGCCTGGCAGCTCTCTA-3′ |
| MX2 | 5′-TGAACGTGCAGCGAGCTT-3′ | 5′-GGCTTGTGGGCCTTAGACAT-3′ |
| PLAT | 5’-AGCGAGCCAAGGTGTTTCAAC-3’ | 5’-TGCCCCTGTAGCTGATGCC-3’ |
| PLAU | 5’-GCTTCTCTGCGTCCTGGTC-3’ | 5’-TGGGCAGTTGCACCAGTGA-3’ |
| TNFSF15 | 5'-CACCACATACCTGCTTGTCAGC-3' | 5'-TCTCCGTCTGCTCTAAGAGGTG-3' |

**Supplemental Table S4: Fluorochrome-labelled antibodies used for flow cytometry**

| **Target** | **Fluorochrome** | **Manufacturer** | **Reference** |
| --- | --- | --- | --- |
| CD3 | BV510 | Biolegend | 317332 |
| CD56 | PE-Cy7 | Beckman Coulter | A21692 |
| IFNγ | FITC | BD Bioscience | 552887 |
| TNF | APC | BD Bioscience | 551384 |
| CD107a | FITC | Biolegend | 328606 |

| Supplemental Table S5. DEG after ruxolitinib withdrawal vs. CHZ868 withdrawal | | | | | | | | | | | | | | | | | | |
| --- | --- | --- | --- | --- | --- | --- | --- | --- | --- | --- | --- | --- | --- | --- | --- | --- | --- | --- |
| Gene_Name | S1DCruxo_Raw.Read | S2-DCruxo_Raw.Read | S3-DCruxo_Raw.Read | S1DCCHZ_Raw.Read | S2DCCHZ_Raw.Read | S3DCCHZ_Raw.Read | S1DCruxo_Normalized | S2DCruxo__Normalized | S3DCruxo_Normalized | S1DCCHZ_Normalized | S2DCCHZ_Normalized | S3DCCHZ_Normalized | baseMean | log2FoldChange | lfcSE | stat | pvalue | padj |
| RGS20 | 1025 | 595 | 721 | 241 | 186 | 171 | 982 | 622 | 721 | 233 | 174 | 179 | 485 | -1.98664094472105 | 0.287212657295507 | -6.91696864416752 | 4.61410162897261e-12 | 7.08864433259063e-08 |
| RFPL4B | 104 | 81 | 95 | 19 | 8 | 9 | 100 | 85 | 95 | 18 | 7 | 9 | 52 | -2.98288436026147 | 0.480902509025193 | -6.20267997001698 | 5.55096481007842e-10 | 3.42214334285477e-06 |
| C3AR1 | 127 | 78 | 79 | 17 | 14 | 13 | 122 | 82 | 79 | 16 | 13 | 14 | 54 | -2.70859434239529 | 0.438750571747184 | -6.17342635386018 | 6.68256852734772e-10 | 3.42214334285477e-06 |
| MX2 | 8413 | 10588 | 6103 | 2947 | 2255 | 2229 | 8061 | 11070 | 6102 | 2855 | 2112 | 2333 | 5422 | -1.78919027337591 | 0.295998811772291 | -6.04458599905569 | 1.49794138080208e-09 | 5.75321835831559e-06 |
| RGS4 | 5224 | 8283 | 4196 | 1554 | 1726 | 1000 | 5005 | 8660 | 4195 | 1505 | 1617 | 1047 | 3672 | -2.09884312440017 | 0.356660590155835 | -5.88470714828103 | 3.98759420639508e-09 | 1.22522819585695e-05 |
| FGF5 | 5586 | 4755 | 4613 | 2313 | 1419 | 1733 | 5352 | 4971 | 4612 | 2241 | 1329 | 1814 | 3387 | -1.4720610414076 | 0.260464861490997 | -5.65166845531861 | 1.58897856449558e-08 | 4.06857961439093e-05 |
| RPL22L1 | 784 | 638 | 872 | 365 | 343 | 336 | 751 | 667 | 872 | 354 | 321 | 352 | 553 | -1.15775693776248 | 0.209546559044143 | -5.52505821638705 | 3.29376376616826e-08 | 7.22887039137757e-05 |
| YPEL2 | 789 | 853 | 770 | 258 | 409 | 328 | 756 | 892 | 770 | 250 | 383 | 343 | 566 | -1.30788467112275 | 0.253186323745858 | -5.16570031024098 | 2.39540337152074e-07 | 0.00037490116853608 |
| CRYBG3 | 2820 | 1860 | 2378 | 1146 | 1205 | 892 | 2702 | 1945 | 2377 | 1110 | 1129 | 934 | 1699 | -1.14654284930255 | 0.222102353382321 | -5.16222737779334 | 2.44028619759214e-07 | 0.00037490116853608 |
| KAT6B | 332 | 336 | 319 | 678 | 646 | 597 | 318 | 351 | 319 | 657 | 605 | 625 | 479 | 0.933186317359861 | 0.180576682187871 | 5.16781184621046 | 2.36850567864606e-07 | 0.00037490116853608 |
| NIPAL1 | 435 | 128 | 234 | 58 | 42 | 25 | 417 | 134 | 234 | 56 | 39 | 26 | 151 | -2.68751464853356 | 0.523139041192437 | -5.13728557212565 | 2.78735226054612e-07 | 0.000389291752534273 |
| CRYBG3 | 1315 | 912 | 1134 | 573 | 603 | 487 | 1260 | 953 | 1134 | 555 | 565 | 510 | 829 | -1.0383263180105 | 0.205064405267981 | -5.06341564570211 | 4.11810644008478e-07 | 0.000527220576991854 |
| IFIT1 | 9729 | 11747 | 4621 | 2802 | 2086 | 1536 | 9322 | 12281 | 4620 | 2714 | 1954 | 1608 | 5417 | -2.06281360571809 | 0.409546508371783 | -5.03682381256071 | 4.73320077080308e-07 | 0.000559355103398829 |
| NET1 | 3769 | 3067 | 3164 | 1942 | 1901 | 1391 | 3611 | 3206 | 3163 | 1881 | 1781 | 1456 | 2517 | -0.963480261047032 | 0.196317588854696 | -4.90776331691884 | 9.2120929586432e-07 | 0.00101089560088311 |
| NCOA7 | 6278 | 4116 | 4918 | 2567 | 2358 | 2556 | 6015 | 4303 | 4917 | 2487 | 2209 | 2675 | 3768 | -1.04757059945568 | 0.218939430408352 | -4.78475072992479 | 1.71199665595304e-06 | 0.00175342697502711 |
| PLAU | 13447 | 22173 | 21052 | 3751 | 6247 | 7176 | 12884 | 23181 | 21047 | 3634 | 5852 | 7511 | 12352 | -1.74854335278979 | 0.367480464258141 | -4.75819403439501 | 1.95332689773662e-06 | 0.0017564043550494 |
| AQP3 | 302 | 175 | 316 | 104 | 86 | 60 | 289 | 183 | 316 | 101 | 81 | 63 | 172 | -1.69039490612091 | 0.356048370388974 | -4.74765522525547 | 2.05788442302214e-06 | 0.0017564043550494 |
| NAV2 | 2256 | 2449 | 2583 | 6156 | 4441 | 6119 | 2162 | 2560 | 2582 | 5964 | 4160 | 6405 | 3972 | 1.17812658753638 | 0.247883086905478 | 4.75275099339723 | 2.00667425194913e-06 | 0.0017564043550494 |
| MAGI3 | 703 | 381 | 531 | 195 | 198 | 233 | 674 | 398 | 531 | 189 | 185 | 244 | 370 | -1.3752062039391 | 0.299668779023636 | -4.5890873531094 | 4.45188115715531e-06 | 0.00359969737986195 |
| CSRP2 | 1434 | 1288 | 1156 | 562 | 811 | 547 | 1374 | 1347 | 1156 | 544 | 760 | 573 | 959 | -1.04621097025734 | 0.231024612626822 | -4.52856930853207 | 5.93844029176066e-06 | 0.00456161291011595 |
| SPTSSA | 1712 | 1508 | 1515 | 934 | 921 | 614 | 1640 | 1577 | 1515 | 905 | 863 | 643 | 1190 | -0.972841246004376 | 0.21851419442037 | -4.45207346179469 | 8.50450584769015e-06 | 0.00622165349228875 |
| TP53INP2 | 3069 | 2303 | 3316 | 1338 | 1343 | 1585 | 2941 | 2408 | 3315 | 1296 | 1258 | 1659 | 2146 | -1.04018554852234 | 0.23575459811763 | -4.41215381090188 | 1.02347361159768e-05 | 0.00708480263265287 |
| CDK17 | 2607 | 2059 | 2285 | 1412 | 1553 | 1196 | 2498 | 2153 | 2284 | 1368 | 1455 | 1252 | 1835 | -0.767185425822611 | 0.174185310592709 | -4.40442091937645 | 1.06066823244819e-05 | 0.00708480263265287 |
| COL3A1 | 54975 | 179834 | 104753 | 17852 | 6843 | 29269 | 52675 | 188013 | 104729 | 17294 | 6410 | 30636 | 66626 | -2.66823485193119 | 0.613017522520115 | -4.35262411580354 | 1.34517671777673e-05 | 0.00826637996608155 |
| DDX58 | 13917 | 19970 | 10278 | 5451 | 6941 | 4460 | 13335 | 20878 | 10276 | 5281 | 6502 | 4668 | 10157 | -1.43521138368789 | 0.329297671879822 | -4.35840124679553 | 1.31016016327752e-05 | 0.00826637996608155 |
| SOX9 | 1431 | 1079 | 1206 | 344 | 658 | 242 | 1371 | 1128 | 1206 | 333 | 616 | 253 | 818 | -1.62234897961688 | 0.374896423823569 | -4.32745920345289 | 1.50839272685071e-05 | 0.0087595554355266 |
| P4HA2 | 10068 | 6900 | 9724 | 5248 | 4091 | 4205 | 9647 | 7214 | 9722 | 5084 | 3832 | 4401 | 6650 | -0.997140752875658 | 0.231090438946586 | -4.31493729217475 | 1.59648214668193e-05 | 0.0087595554355266 |
| ERF | 249 | 232 | 257 | 425 | 471 | 467 | 239 | 243 | 257 | 412 | 441 | 489 | 347 | 0.862032765223422 | 0.199766945653545 | 4.31519219760432 | 1.59464101649204e-05 | 0.0087595554355266 |
| GATA6 | 1476 | 2227 | 1440 | 557 | 902 | 608 | 1414 | 2328 | 1440 | 540 | 845 | 636 | 1201 | -1.35830088384287 | 0.318830239037583 | -4.26026366866274 | 2.0418588349217e-05 | 0.0108169232003111 |
| BCL2L14 | 39 | 44 | 52 | 13 | 9 | 6 | 37 | 46 | 52 | 13 | 8 | 6 | 27 | -2.30470495224927 | 0.546791871721129 | -4.21495832590704 | 2.49824316452529e-05 | 0.012793503245534 |
| TMEM2 | 22412 | 12001 | 15554 | 8976 | 6938 | 5524 | 21474 | 12547 | 15550 | 8695 | 6499 | 5782 | 11758 | -1.24070763195324 | 0.300170800025768 | -4.13333885856564 | 3.57530995069992e-05 | 0.0163635016940976 |
| CPNE8 | 1618 | 943 | 1164 | 677 | 600 | 462 | 1550 | 986 | 1164 | 656 | 562 | 484 | 900 | -1.12066399648317 | 0.271321337080152 | -4.13039390319721 | 3.62142197226659e-05 | 0.0163635016940976 |
| TRIM5 | 2273 | 2759 | 2533 | 1376 | 1605 | 1391 | 2178 | 2884 | 2532 | 1333 | 1504 | 1456 | 1981 | -0.823102882409218 | 0.198586413511697 | -4.14480964661127 | 3.40096107016487e-05 | 0.0163635016940976 |
| DCAF5 | 3957 | 4230 | 3776 | 2651 | 2613 | 2330 | 3791 | 4422 | 3775 | 2568 | 2448 | 2439 | 3241 | -0.685410369811941 | 0.165681241808699 | -4.13692197335977 | 3.51995736229269e-05 | 0.0163635016940976 |
| SPSB2 | 31 | 50 | 37 | 97 | 135 | 122 | 30 | 52 | 37 | 94 | 126 | 128 | 78 | 1.55155520600643 | 0.379359085201908 | 4.08993817870643 | 4.31488182199377e-05 | 0.0189398655517972 |
| SEC24D | 10776 | 9994 | 9666 | 7164 | 6909 | 6527 | 10325 | 10449 | 9664 | 6940 | 6472 | 6832 | 8447 | -0.588344094607459 | 0.144304844503307 | -4.07709177493329 | 4.56024967423079e-05 | 0.0194608654847799 |
| ISG20 | 671 | 456 | 473 | 213 | 300 | 258 | 643 | 477 | 473 | 206 | 281 | 270 | 392 | -1.07246808399668 | 0.264805301626526 | -4.05002497083406 | 5.12121679057541e-05 | 0.0212641225820568 |
| BRMS1L | 588 | 491 | 459 | 296 | 317 | 284 | 563 | 513 | 459 | 287 | 297 | 297 | 403 | -0.80182007270095 | 0.198515693301245 | -4.03907650507105 | 5.36620615921044e-05 | 0.0215593717796785 |
| CCDC109B | 1561 | 1297 | 1489 | 1000 | 935 | 792 | 1496 | 1356 | 1489 | 969 | 876 | 829 | 1169 | -0.698948353224455 | 0.173333058314073 | -4.03240074353265 | 5.52099357478056e-05 | 0.0215593717796785 |
| KLF10 | 2160 | 2592 | 2011 | 4549 | 4099 | 3830 | 2070 | 2710 | 2011 | 4407 | 3840 | 4009 | 3174 | 0.852030511582296 | 0.211500547623872 | 4.02850262637395 | 5.61332338206821e-05 | 0.0215593717796785 |
| PLEKHA4 | 2652 | 3149 | 2140 | 1281 | 1269 | 1440 | 2541 | 3292 | 2140 | 1241 | 1189 | 1507 | 1985 | -1.01804370153983 | 0.253142307891541 | -4.02162605697666 | 5.77977629746973e-05 | 0.0216572446970799 |
| LEPRE1 | 3903 | 4394 | 4025 | 2591 | 2507 | 2543 | 3740 | 4594 | 4024 | 2510 | 2349 | 2662 | 3313 | -0.716541877449495 | 0.179275093377103 | -3.99688469798903 | 6.41815424416601e-05 | 0.0234766913459815 |
| SLC30A7 | 5720 | 4770 | 4809 | 3362 | 3564 | 2645 | 5481 | 4987 | 4808 | 3257 | 3339 | 2769 | 4107 | -0.705938250382288 | 0.178110779795744 | -3.96347852270285 | 7.38655117155507e-05 | 0.026390601313628 |
| LRRC8C | 4066 | 1808 | 2087 | 1212 | 1015 | 673 | 3896 | 1890 | 2087 | 1174 | 951 | 704 | 1784 | -1.47631095076637 | 0.373535630107796 | -3.95226273418772 | 7.7415673967907e-05 | 0.0266798724984064 |
| ZSWIM6 | 3542 | 1794 | 2967 | 1386 | 1305 | 1179 | 3394 | 1876 | 2966 | 1343 | 1223 | 1234 | 2006 | -1.116242597406 | 0.282915630139281 | -3.94549639005972 | 7.96348318642068e-05 | 0.0266798724984064 |
| XPO1 | 4579 | 4510 | 4754 | 7319 | 7010 | 6936 | 4387 | 4715 | 4753 | 7090 | 6567 | 7260 | 5795 | 0.594229716831929 | 0.150638322380292 | 3.94474465356681 | 7.98850572757075e-05 | 0.0266798724984064 |
| BLZF1 | 3338 | 3321 | 2678 | 1972 | 1951 | 1453 | 3198 | 3472 | 2677 | 1910 | 1828 | 1521 | 2434 | -0.829737762620723 | 0.210779385206512 | -3.93652235871066 | 8.26709210749796e-05 | 0.027022837456913 |
| DUSP6 | 8813 | 9028 | 5663 | 2352 | 4504 | 1830 | 8444 | 9439 | 5662 | 2278 | 4219 | 1915 | 5326 | -1.48458383868178 | 0.382268221542634 | -3.88361824242354 | 0.000102913448771942 | 0.0322665166017009 |
| SLC20A1 | 10622 | 8138 | 8716 | 5719 | 4784 | 3054 | 10178 | 8508 | 8714 | 5540 | 4482 | 3197 | 6770 | -1.05156448416454 | 0.270707375154852 | -3.88450622582047 | 0.000102538065442567 | 0.0322665166017009 |
| LMO2 | 249 | 227 | 370 | 104 | 53 | 116 | 239 | 237 | 370 | 101 | 50 | 121 | 186 | -1.63936201746043 | 0.423288178169845 | -3.87292181073538 | 0.00010753833541439 | 0.0326799709418132 |
| MMP19 | 1297 | 1173 | 1057 | 787 | 605 | 625 | 1243 | 1226 | 1057 | 762 | 567 | 654 | 918 | -0.830111903516989 | 0.214455834236936 | -3.87078256215618 | 0.000108486527242887 | 0.0326799709418132 |
| RASL11A | 164 | 194 | 182 | 83 | 105 | 66 | 157 | 203 | 182 | 80 | 98 | 69 | 132 | -1.12641802194998 | 0.292826538527535 | -3.84670743169019 | 0.000119715780186172 | 0.0353691063653877 |
| C4orf32 | 1202 | 1853 | 1129 | 705 | 701 | 566 | 1152 | 1937 | 1129 | 683 | 657 | 592 | 1025 | -1.12608356163995 | 0.29547797128152 | -3.81105757818764 | 0.000138373529491587 | 0.0366522850617113 |
| SPRED1 | 10820 | 10203 | 7873 | 5247 | 5504 | 2918 | 10367 | 10667 | 7871 | 5083 | 5156 | 3054 | 7033 | -1.12058319392872 | 0.29356305813749 | -3.81718054389492 | 0.000134985394720745 | 0.0366522850617113 |
| ELOVL6 | 2286 | 1963 | 2009 | 900 | 1470 | 878 | 2190 | 2052 | 2009 | 872 | 1377 | 919 | 1570 | -0.980394814342323 | 0.256964099022563 | -3.81529878326014 | 0.000136018254241204 | 0.0366522850617113 |
| SCYL2 | 4009 | 3196 | 3696 | 2560 | 2215 | 2167 | 3841 | 3341 | 3695 | 2480 | 2075 | 2268 | 2950 | -0.672924319320111 | 0.175560638946874 | -3.83300222280315 | 0.000126588809220043 | 0.0366522850617113 |
| GBP5 | 4879 | 6668 | 5706 | 9477 | 15260 | 11885 | 4675 | 6971 | 5705 | 9181 | 14295 | 12440 | 8878 | 1.04966789503898 | 0.27534590438483 | 3.81217907484087 | 0.000137747018153568 | 0.0366522850617113 |
| BRICD5 | 36 | 28 | 28 | 77 | 78 | 107 | 34 | 29 | 28 | 75 | 73 | 112 | 59 | 1.49818798283006 | 0.392021855775751 | 3.82169504265362 | 0.000132537527555241 | 0.0366522850617113 |
| SMAP2 | 1660 | 1642 | 2088 | 1096 | 761 | 943 | 1591 | 1717 | 2088 | 1062 | 713 | 987 | 1359 | -0.966132199055545 | 0.253825581515006 | -3.80628380043099 | 0.00014107049839741 | 0.0367333231674477 |
| TPST1 | 2634 | 2085 | 2125 | 1363 | 1594 | 1062 | 2524 | 2180 | 2125 | 1320 | 1493 | 1112 | 1792 | -0.798563898725708 | 0.21027119287518 | -3.79778079824633 | 0.000145997344057355 | 0.0373826199458857 |
| RCOR3 | 331 | 294 | 356 | 664 | 528 | 694 | 317 | 307 | 356 | 643 | 495 | 726 | 474 | 0.92671339398267 | 0.244349756726658 | 3.79256933338935 | 0.000149096603466757 | 0.037550346214095 |
| SLC37A3 | 6583 | 5867 | 5254 | 3901 | 4196 | 3348 | 6308 | 6134 | 5253 | 3779 | 3931 | 3504 | 4818 | -0.657911053100257 | 0.173669912519833 | -3.7882845885877 | 0.000151691033553009 | 0.0375875701366916 |
| MIDN | 670 | 537 | 912 | 1199 | 2090 | 1689 | 642 | 561 | 912 | 1162 | 1958 | 1768 | 1167 | 1.20820885706641 | 0.31956040054821 | 3.78084660988566 | 0.000156295953943922 | 0.0381138847688963 |
| RP11-39K24.14 | 42 | 43 | 30 | 12 | 8 | 5 | 40 | 45 | 30 | 12 | 7 | 5 | 23 | -2.23668354823066 | 0.592271633613798 | -3.77644888137447 | 0.000159080189622913 | 0.0381867023933878 |
| NCR3LG1 | 659 | 453 | 580 | 358 | 272 | 269 | 631 | 474 | 580 | 347 | 255 | 282 | 428 | -0.932152617902822 | 0.248485896692091 | -3.7513300767242 | 0.000175898947341786 | 0.0415743927386439 |
| RP11-443P15.2 | 162 | 267 | 172 | 424 | 475 | 520 | 155 | 279 | 172 | 411 | 445 | 544 | 334 | 1.20791425341919 | 0.323238794603525 | 3.73690990557238 | 0.000186295637110094 | 0.0420891157782702 |
| IDO2 | 0 | 5 | 3 | 25 | 39 | 25 | 0 | 5 | 3 | 24 | 37 | 26 | 16 | 3.42122639925899 | 0.91395618410448 | 3.74331555359101 | 0.000181607925089245 | 0.0420891157782702 |
| SPOCK2 | 2 | 6 | 3 | 13 | 38 | 68 | 2 | 6 | 3 | 13 | 36 | 71 | 22 | 3.42287774091106 | 0.915629955889689 | 3.73827627514126 | 0.00018528627237936 | 0.0420891157782702 |
| ATF3 | 7702 | 5945 | 6508 | 4585 | 4265 | 4278 | 7380 | 6215 | 6507 | 4442 | 3995 | 4478 | 5503 | -0.638327324721081 | 0.171125104219717 | -3.73017931899401 | 0.000191343540376377 | 0.0426030552290186 |
| MGP | 33 | 264 | 43 | 10 | 17 | 12 | 32 | 276 | 43 | 10 | 16 | 13 | 65 | -3.19787566896708 | 0.859817391801997 | -3.71924980752599 | 0.000199815359987708 | 0.0438537625070165 |

**Supplemental Methods**

*In Vitro Studies*

For the described experiments, MSCs were expanded in α-minimal essential media (αMEM) (Corning, Tewksbury, MA) and 10% fetal bovine serum (FBS) (Sigma-Aldrich, St. Louis, MO) supplemented with 1% penicillin/streptomycin (Lonza, Walkersville, MD) and L-glutamine (Corning) and grown to 80% confluence. Primary pooled human umbilical vein endothelial cells (HUVECs) (cat. # PCS-100-013, ATCC, Manassas, VA) were cultured at starting seeding density of 3,000-5,000 cells/cm^2^ in accordance with the manufacturer’s recommendations using vascular cell basal medium supplemented with bovine brain extract (cat. # PCS-100-040, ATCC, Manassas, VA) and passaged three times before use in experiments. Optimal concentrations of IFNγ and drugs were determined in preliminary experiments.

MSCs and HUVECs were treated with the following reagents in the described experiments: IFNγ (10 ng/mL), ruxolitinib (1 µM unless otherwise specified), baricitinib (1 µM unless otherwise specified), CHZ868 (1 µM unless otherwise specified). Next, they were washed with PBS and then treated with regular growth media with 0.01% DMSO or 10 ng/mL final concentration IFNγ+0.01% DMSO or 10 ng/mL IFNγ+drug for 48 hours. In the event of withdrawal, the cells were washed twice and then treated with our usual growth media with either 0.01% DMSO or with 10 ng/mL IFNγ+0.01% DMSO for a variety of time periods before harvest. In all cases the harvest of the cells was performed by first collecting the conditioned media, then washing once with PBS on ice, then removal all traces of PBS and conducting on-plate lysis (cat. #9803, Cell Signaling Technology, Danvers, MA) supplemented with 1 mM EDTA, 1 mM PMSF, 1 mM NaF and 1 mM Na_3_VO_4_.

*Western Blot*

We loaded equal amounts of protein from lysates of SG-MSC or HUVECs to run on a tris-glycine-SDS 10% polyacrylamide gel (cat. #161-0374, Bio-Rad, Hercules, CA). This electrophoresis, in tris-glycine-SDS buffer at room temperature, was at 50V for ~45 minutes followed by ~135 minutes at ~90 V. Next, after incubating the gels on ice for 10 minutes, a ‘sandwich’ in a gel holder cassette – containing mesh, filter paper, the gel, the nitrocellulose membrane (0.2 µm), filter paper and mesh, from cathode to anode – was made for transfer of proteins from each gel to each membrane (cat. #1620112, BioRad, Hercules, CA) at 90 V for 90 minutes, with the chamber surrounded by ice in the cold room. The transfer buffer was tris-glycine-SDS buffer with the addition of 20% methanol. The transfer was verified by rinsing the membranes with water briefly, then incubating them with 0.1% Ponceau S (cat. #141194, Millipore Sigma, St. Louis, MO) in 5% aqueous acetic acid, then rinsing twice in water. Next, the membranes were washed twice with tris-buffered saline (20 mM tris, 150 mM NaCl, pH 7.6) + 0.1% Tween20 (TBST).

The membranes were blocked with 5% non-fat dry milk (cat. #sc-2325, Santa Cruz Biotechnology, Dallas, TX) in TBST (for phospho and total TYK2, 3% BSA in TBST was used per manufacturer instructions) at room temperature for ~ 1 hr. Primary antibodies were diluted 1:1,000 (typically) in either 5% milk in TBST+0.02% sodium azide or 5% BSA in TBST+0.02% sodium azide, and their incubation with membranes was performed overnight at 4° C with gentle rocking. Next, three washes at room temperature were performed with TBST, ~5-10 minutes per wash. This was followed by a ~2 hr incubation with gentle rocking at room temperature with HRP-conjugated secondary antibodies raised in goat against primary antibodies from rabbit or mouse (cat. #A120-101P, and #A90-116P, respectively, Bethyl Laboratories, Montgomery, TX) diluted 1:5,000 in 5% milk in TBST. Subsequently, three washes with TBST were performed at room temperature, ~5-10 minutes per wash. The membranes wer placed face-down in a 1:1 mixture of the luminol-enhancer and peroxide components of Clarity™ Western ECL Substrate (cat.# 1705060, BioRad, Hercules, CA). After the incubation, the membranes were imaged Amersham ImageQuant 800 (Cytiva, Marlborough, MA). Densitometric analysis was performed using ImageStudio Lite software (Li-Cor, Lincoln, NE).

Primary antibodies used were principally from Cell Signaling Technology: phospho-Akt (#4060), total Akt (#9272), phospho-Erk (#4370), total Erk (#9102), phospho-JAK1 (#3331), total JAK1 (#3332), phospho-JAK2 (#3771), total JAK2 (#3230), phospho-STAT1 (cat. # 9167), total STAT1 (cat. #9172), phospho-STAT2 (#88410), phospho-STAT3 (#9145), total STAT3 (#9139), total STAT4 (#2653), phospho-STAT5 (#9359), total STAT5 (#95205); others were from Abcam: total STAT2 (#32367), phospho-STAT4 (#ab313630); Bioss: phospho-TYK2 (#bs3437R), total TYK2 (#bs6662R); and Santa Cruz Biotechnology: GAPDH (sc-25778).

*RNA Seq*

Four each of three sicca control MSC lines, we grew cells as described above up until the withdrawal time of 3 hours was over. At that point, we collected conditioned media, washed the cells twice with PBS supplemented with 10 ng/mL IFNγ+DMSO or 10 ng/mL IFNγ+1μM ruxolitinib for the condition with IFNγ+1μM ruxolitinib throughout. ew added 5 mL of 0.05%trypsin/0.5mM EDTA supplemented similarly and detached cells for 5 min in the incubator. Double the volume (10 mL) of growth media supplemented by 10 ng/mL IFNγ with or without 1μM ruxolitinib was added for neutralization of trypsin, and the cells were collected by centrifugation at 1,500 rpm for 5 min. After discarding the supernatant, the cell pellet was resuspended in 1 mL of PBS supplemented by 10 ng/mL IFNγ with or without 1μM ruxolitinib, transferred to a microcentrifuge tube and spun at ~1,000x*g* for 5 min at 4° C. Next, the PBS supernatant was discarded, and the cell pellet was resuspended in 1 mL sterile cryostor. The microcentrifuge tubes were placed at 4$^{\circ}$ C for ~ 10 min in a controlled cooling system and then transferred to -80$^{\circ}$ C overnight with controlled cooling. The tubes were placed and maintained at -80$^{\circ}$ C until shipment to MedGenome, Inc. (Foster City, CA) for RNA isolation, reverse transcription, library construction and sequencing. The read alighnment was performed with STAR (v2.7.3a) aligner. Reads mapping to ribosomal and mitochondrial genes were removed before performing aligntment. Raw read counts were estimated with HTSeq (v0.11.2) and normalized with DESeq2. The alignedd reads were used to estimate expression of genes with cufflinks (v2.2.1). Distribution of mappe dreads were performed with RSeQC and RNA-SeQC. Hierarchical clustering, heatmap and correlation analysis was performed with R (v3.5.2) and additional associated packages. Differential expression analysis was performed with DESeq2 (R Bioconductor package).

*Real time quantitative PCR*

After harvesting the MSCs in each treatment condition in TRIzol^®^, we used the Direct-zol RNA Miniprep columns for RNA isolation (cat. # Zymo Research, Irvine, CA). We generated cDNA using SuperScript IV reverse transcriptase (cat. #18090050, Invitrogen, Waltham, MA) by combining 750 ng of isolated RNA for each sample with RNAse free water to reach 11 µL, and then adding 1 µL of 50 µM oligo-dT_22_ primer (Integrated DNA Technologies, Coralville, IA) and 1 µL of a mixture having 10 mM of each dNTP (diluted into water from cat. #10297, Invitrogen, Waltham, MA). These 13 µL reactions were placed at 65° C for 5 minutes, and then on ice for ~ 3 min and then spun down. Next, each reaction received 4 µL of the manufacturer’s 5x SSIV buffer, 1 µL of 100 mM DTT, 1 µL of SuperScript IV enzyme solution and 1 µL of RNAseOut (cat. #10777019, Invitrogen, Waltham, MA). After mixing well and spinning down, the now-20 µL reactions were placed at 52° C for 10 min, left in the heat block as the temperature rose to 80° C over ~ 2 min, and left at 80° C for 10 min more. The reactions were then set on ice, spun down and stored at -20° C until qPCR was run. The qPCR was performed using the QuantiNova SYBR Green kit (*PLAU, PLAT*) (cat. # 208052, Qiagen, Germantown, MD) or the qPCR 2x Green Master Mix (*IFIT1*, *MX1*, *MX2* and *TNFSF15*) (cat. # 42-116PG, Apex Reagents and Chemicals). Primers sequences are shown in Table S3. ΔΔC_t_ values (ΔΔC_t_=ΔC_t,IFNγ all the way_-ΔC_t,__) are plotted in the figure, and ΔC_t_ values were used for determining statistical significance.

*ELISA*

Enzyme-linked immunosorbent assay (ELISA) was performed on conditioned media (CM) collected above the adherent cells. Cells were grown to 80% confluence before collection of medium. This CM was spun at 4,000 rpm for 10 min at 4° C to remove adventitious whole cells and decanted to fresh tubes. Aliquots were then stored at -80° C. They were thawed on ice and mixed well before beginning assays. The uPA ELISA kits were performed per the manufacturer’s instructions (for the MSC experiment, cat. #IHUUPAKTT, Innovative Research, Novi, MI; for the HUVEC experiment, cat. #CSB-E04751h, CUSABIO, Houston, TX).
